# Computational analysis of SOD1-G93A mouse muscle biomarkers for comprehensive assessment of ALS progression

**DOI:** 10.1101/2024.03.11.584407

**Authors:** Pedro Gómez-Gálvez, Victoria Navarro, Ana M. Castro, Carmen Paradas, Luis M. Escudero

## Abstract

**Aims:** To identify potential image biomarkers of neuromuscular disease by analysing morphological and network-derived features in skeletal muscle biopsies from a murine model of amyotrophic lateral sclerosis (ALS), the SOD1^G93A^ mouse, and wild-type (WT) controls at distinct stages of disease progression.

**Methods:** Using the NDICIA computational framework, we quantitatively evaluated histological differences between skeletal muscle biopsies from SOD1^G93A^ and WT mice. The process involved the selection of a subset of features revealing these differences. A subset of discriminative features was selected to characterise these differences, and their temporal dynamics were assessed across disease stages.

**Results:** Our findings demonstrated that muscle pathology in the mutant model evolves from early alterations in muscle fibre arrangement, detectable at the presymptomatic stage through graph theory features, to the subsequent development of the typical morphological pattern of neurogenic atrophy at more advanced disease stages.

**Conclusions:** Our assay identifies a neurogenic signature in mutant muscle biopsies, even when the disease was phenotypically imperceptible.

**KEY POINTS:**

- NDICIA analysis detected differences between SOD1^G93A^ and WT muscles at presymptomatic stage through the analysis of graph theory features.
- Our computational tool identified different neurogenic-like traits in the ALS mouse model at all analysed stages of disease progression.
- Differences between SOD1^G93A^ and WT muscle images became more pronounced as the disease advanced.
- The integration of UMAP into the upgraded NDICIA framework was validated as a robust alternative to PCA.
- Muscle fibre characteristics in SOD1^WT^ closely resembled those of WT mice.

## INTRODUCTION

Amyotrophic Lateral Sclerosis (ALS) is the most common neurodegenerative disorder affecting the motor neuron system [1,2]. The lack of effective treatments for this devastating disease [2,3] highlights the urgency of new therapeutic approaches. Most ALS cases are sporadic [2,4], but approximately 10% are familial and linked to mutations in genes, including *C9ORF72*, *SOD1*, *FUS* and *TARDBP* (TDP-43) [5–8].

Mouse models have been critical in ALS research, providing valuable insights into disease mechanisms and therapeutic strategies [9,10]. Among murine models, SOD1^G93A^ mice are the most extensively studied due to their robust and reproducible phenotype, which mirrors key aspects of human ALS, such as progressive motor dysfunction (Gurney, 1997; Gurney et al., 1994; Weydt et al., 2003). Recent research indicates a wide variability in the disease onset times, depending on the mice strain [13]. The model used in this work reports a disease onset around 90 days of age [14,15], with rapid progression leading to death around the 20th week [16,17].

The examination of skeletal muscle biopsies offers a promising avenue for uncovering biomarkers and understanding the mechanisms underlying ALS pathology. Skeletal muscle is composed of tightly packed fibres forming a mosaic-like arrangement connected by endomysium. This organisation reflects physiological constraints and is influenced by the metabolic properties of muscle fibres, which can be classified as type I (slow) or type II (fast) based on the expression of different myosin isoforms [18,19]. Changes in muscle fibre morphology, collagen network density, arrangement and fibre type distribution are hallmarks of neurogenic diseases, including ALS [20–22].

Advances in the computational analysis of images have facilitated the extraction of structural and morphological information from muscle tissues and their objective analysis [22–25]. NDICIA (Neuromuscular DIseases Computerized Image Analysis) [24], developed and patented in our laboratory, is an objective and reproducible method for analysing tissue organisation. It considers the tissue as a graph, where fibres are represented as nodes and their spatial relationships as edges. From its foundation, NDICIA has used PCA (Principal Component Analysis) to reduce the dimensionality of image features, enabling an easier analysis of how much two groups of images differ, and a filtering of the most relevant features associated with the differentiation. However, algorithms that incorporate non-linear transformations to reduce dimensionality, such as UMAP (Uniform Manifold Approximation and Projection [26]), have emerged as promising to enhance or complement NDICIA outcomes.

In this study, we used NDICIA pipelines with PCA and UMAP methods to analyse the skeletal muscle organisation in SOD1^G93A^ mice. Our analysis identified significant differences in the organisational patterns between pre-symptomatic and late stages of the disease, providing a neurogenic signature of the mutant condition.

## MATERIAL AND METHODS

### Transgenic mice

The SOD1^G93A^ transgenic mice colony was obtained by crossing B6SJL-Tg(SOD1-G93A)1Gur/J [JAX #002726] males with wild-type B6SJL (WT) females to maintain the genetic background. This transgenic strain, known as the G1H line, carries approximately 25 copies of the transgene, leading to an aggressive disease course with signs of motor impairment appearing at around 90 days of age [15]. Non-transgenic littermates were used as age-matched controls (WT). The SOD1^WT^ mice colony, carrying the normal allele of the human *SOD1* gene, was maintained hemizygous by crossing B6SJL-Tg(SOD1)2Gur/J [JAX #002297] males with wild-type B6SJL females, maintaining the genetic background [12]. All animals were provided by the Jackson Laboratories. All animal experiments were performed according to the Spanish and European Union regulations (RD53/2013 and 2010/63/UE) and approved by the Animal Research Committee from the Hospital Virgen del Rocío (Seville, Spain). Animals were housed under controlled temperature and humidity conditions, alternating 12-hour light cycles. Experiments and procedures with animals were designed to minimise animal suffering and reduce the number of animals used.

### Tissue sampling

Samples were obtained from SOD1^G93A^ and WT mice at three different ages established to track the progression of ALS disease: a presymptomatic group, at postnatal day 60 (p60), where the mice are mostly non-symptomatic for ALS; an intermediate group, at 100 days (p100), represents a progressing disease stage; and the last group at 120 days of age (p120), where most of the mice are severely affected by motor symptoms. SOD1^WT^ mice muscle was only imaged at p120 to be compared against WT and SOD1^G93A^ mice.

Tissue samples were obtained from each mouse line at the corresponding time point. To extract muscle samples, animals were euthanised via cervical dislocation, and the soleus muscle was isolated from the hindlimbs. Muscle tissue was preserved by snap freezing. To avoid damage to the cells during the process, the whole muscle was first covered completely using talcum powder and then immersed in liquid nitrogen for 1 minute. All samples were placed in cryogenic safe tubes and stored at −80°C. The number and sex of animals varied based on their experimental availability. For each animal, we took the maximum number of images from non-overlapping regions free from artefacts (e.g., tissue breaks, gaps, borders). Information regarding the number of mice, the number of images per group and genotype, mouse sex, left/right soleus, and specific details on image and mouse counts are provided in **Table S1**.

### Immunofluorescence

For immunofluorescence, the soleus was cut transversally into 10 µm sections, using a cryostat (Leica), and placed sequentially onto Superfrost™ Plus microscope slides, to obtain serial slices of each muscle sample. Tissue slices were fixed using 4% paraformaldehyde for 20 minutes at room temperature and permeabilised by incubating for 10 minutes with 0.1% Triton X-100 in PBS. The tissue was then blocked using 1% BSA for 1 hour at room temperature and incubated with primary antibodies overnight at 4°C. The following antibodies were used: to detect muscle fibre types we used mouse anti-myosin heavy chain (slow) (Leica, clone WB-MHCs; 1:200); to delimit the fibres we used rabbit anti-collagen type VI (Abcam, ab6588, 1:300). Finally, slices were incubated for 1 hour at room temperature with the following secondary antibodies: Cy3 donkey anti-mouse (#715-165-151, Jackson Immunoresearch, 1:500) or Alexa488 donkey anti-rabbit (#715-545-150, Jackson Immunoresearch, 1:500). After staining, the preparations were mounted using Fluoromount-G (#00-4958-02, Invitrogen). All images were captured with a 40X objective, under identical conditions and using the same settings in a fluorescence microscope (Nikon Eclipse Ti-E), with an XY resolution of 0.104 µm per pixel, and an image size of 4080 x 3072 pixels.

### NDICIA pipeline

NDICIA is structured as a modular workflow that takes two subsets of muscle tissue images and then processes, analyses and compares them for a predefined set of features. The set of features to be analysed can be customised based on the specific objectives of the study. In this work, we utilised 67 features, subdivided into pure geometric features [10 ccs], pure geometric adding the influence of slow fibre discrimination [15 ccs], and all the features [67 ccs] (**Table S2**). These subsets allowed us to evaluate the contribution of morphological, slow-fibre-related, and graph theory-based features.

NDICIA objectively evaluates differences between groups of images and identifies those features driving group separation. The general workflow is: 1) Image pre-processing, which involves noise reduction and intensity normalisation ensuring image quality optimised for segmentation; 2) Image segmentation, which isolates key components of muscle tissue, such as muscle fibres and collagen network (**Figure 2E**), using a combination of intensity thresholding and morphological operation; 3) Slow fibre identification, which classifies muscle fibres into slow and non-slow categories based on intensity thresholds applied to the red channel (myosin heavy chain [slow] staining, **Figure 2B**); 4) Feature extraction module, which captures various geometric and network features from the segmented images (**Table S2**); and 5) Feature selection module, performs dimensionality reduction (PCA or UMAP) on the extracted features and selects the ones which maximise separation among image classes, based on a cluster separation descriptor. A separation descriptor value greater than 1 indicates good inter-class differentiation, with higher values reflecting more pronounced differences between image classes. The output is the combination of features providing the highest separation descriptor, the absolute value of the descriptor and a plot where each image is projected as a dot in a 2D space through the reduction of dimensionality of the chosen features (**Figures 3**, **4** and **5B-E**). For technical details on the NDICIA pipeline, see the **Supplementary Methods**.

**Figure 1.**
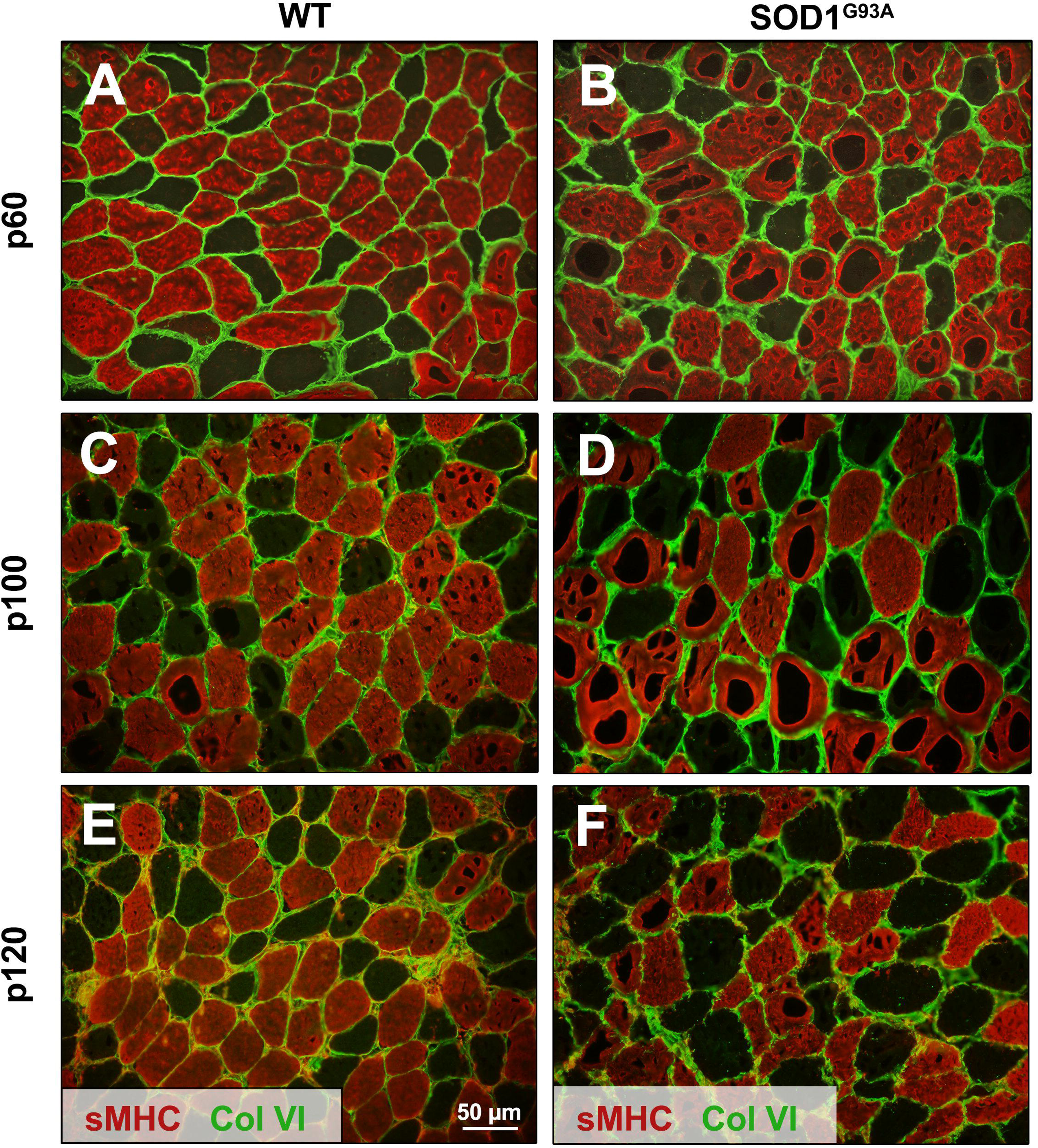
Images of WT and SOD1^G93A^ soleus muscles biopsies. Fluorescence images showing slow (sMHC, red stained) and non-slow fibres (non-stained, observable in dark green as the counterpart of slow fibres), connected by collagen VI content, including the endomysium and perimysium (light green). Collagen demarcates the fibres’ outlines, and slow fibres are indicated by slow myosin heavy chain staining. (**A**, **C**, **E**) Images from WT samples of the soleus muscle. (**B**, **D**, **F**) Images from SOD1^G93A^ mutant mice. The images are also classified by age group: p60 (**A**, **B**), p100 (**C**, **D**) and p120 (**E**, **F**). sMHC tags for myosin heavy chain in slow fibres, while col VI indicates the collagen VI staining. Some cryoartefacts are visible within muscle fibres in the red channel. The scale bar represents 50 µm.

**Figure 2.**
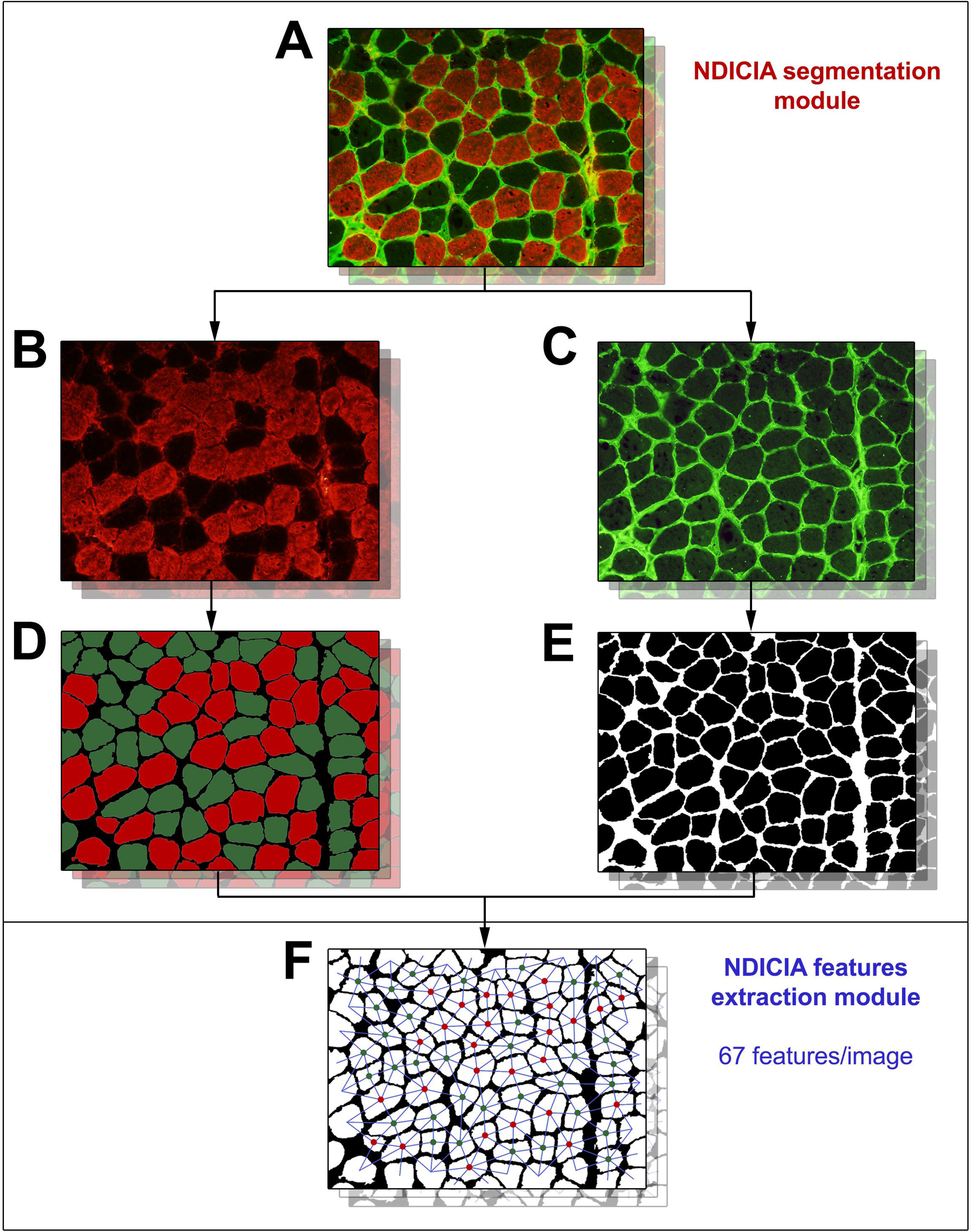
NDICIA features an extraction pipeline. NDICIA processes the images from the microscope (**A**), leveraging the information provided by the red channel (**B**), staining the slow muscle fibres, and the green channel (**C**), staining the collagen VI that delimits the cell compartments. The green channel allows for the segmentation of muscle fibres (**C**, **E**), while the red channel is for fibre type detection (**B**, **D**). Cryoartefacts do not alter or disturb fibre segmentation. A connectivity network is built by linking neighbouring muscle fibres (**F**), and a set of 67 characteristics (geometric, network and fibre type proportion) are extracted. In (**D**), red segmented cells represent slow fibres, while the dark green ones represent non-slow fibres. In (**F**), the same colour code applies to the nodes used to construct the tissue network that discriminates the type of fibre, where the neighbouring cells are linked by a blue line.

**Figure 3.**
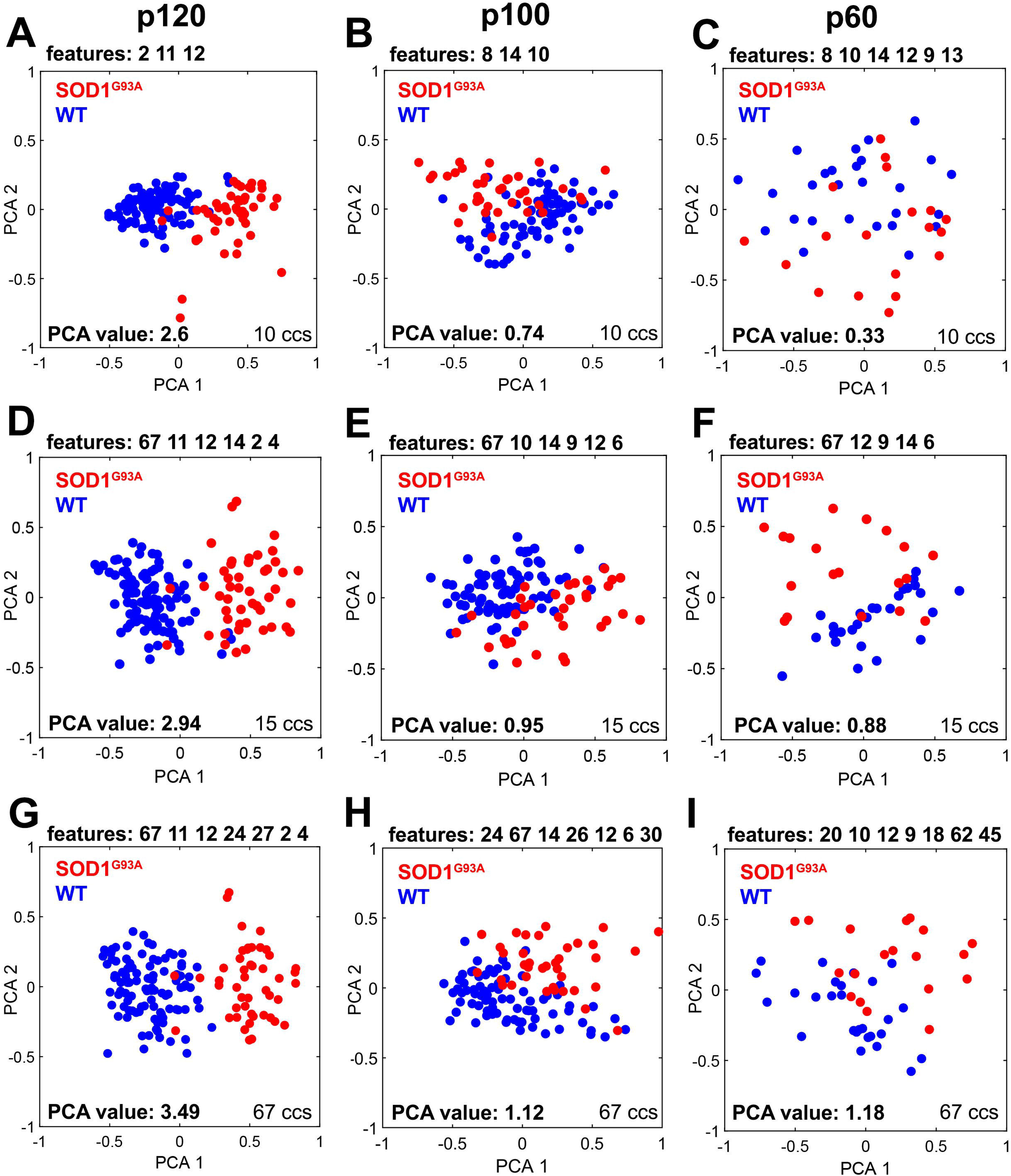
WT vs SOD1^G93A^, NDICIA comparison (PCA). PCA graphs were generated as outputs by NDICIA after feature selection. The selected characteristics are labelled with their numerical index above each graph and arranged from highest to lowest weight in separation, left to right. Each illustrates the reduction of dimensionality from the selected features to the two principal components (PCA 1 on the X-axis and PCA 2 on the Y-axis). These graphs are categorised into three groups based on the subset used as the source for NDICIA: Geometric (10 ccs, **A, B, C**), Geometric including slow fibre discrimination (15 ccs, **D, E, F**) and the full set of features (67 ccs, **G, H, I**). Furthermore, the PCA graphs are further subdivided by age: p120 (**A, D, G**), p100 (**B, E, H**) and p60 (**C, F, I**). Red dots represent each SOD1^G93A^ sample, while blue dots the WT ones. The PCA values plotted on the graphs show how distinct the SOD1^G93A^ and WT groups are: the closer the value is to 0, the more they overlap, while values greater than 1 indicate a clear separation between the groups (**Material and Methods**).

**Figure 4.**
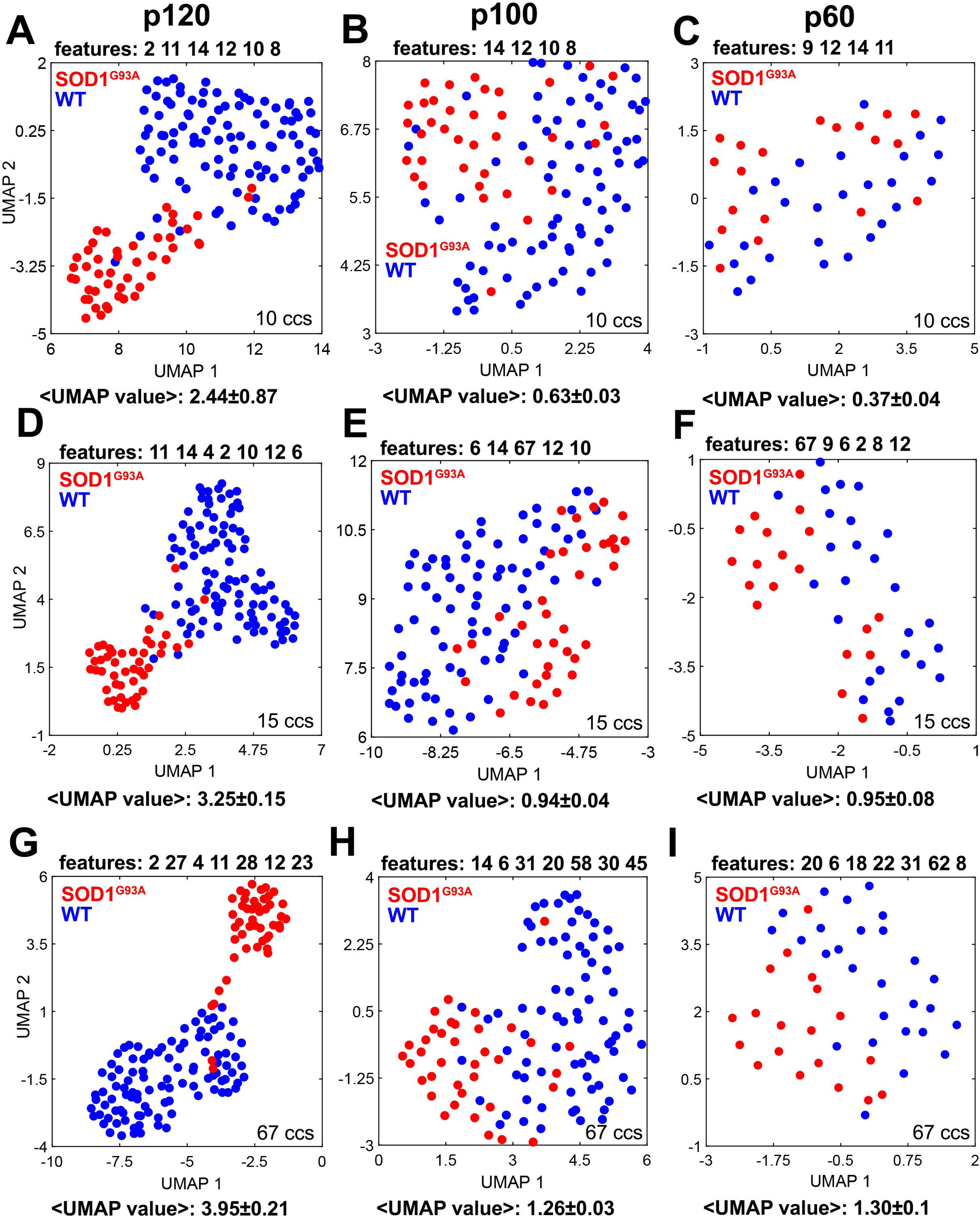
WT vs SOD1^G93A^, NDICIA comparison (UMAP). Similar panels as in Figure 3 but using UMAP as a reduction of dimensionality method instead of PCA. The selected features, shown above the graphs from left to right, are the most frequently chosen across 100 repetitions of the NDICIA pipeline using UMAP. The number of features, with a maximum of 7 features as in PCA, is determined by calculating the mode value after evaluating the number of selected features across 100 randomizations (**Material and Methods**). The UMAP value displays the average and standard deviation considering the 100 repetitions (**Material and Methods**). Additionally, each graph displayed in the different panels embodies the randomization with the more similar UMAP value to the average UMAP value. The colour code and panel order are the same as in Figure 3.

**Figure 5.**
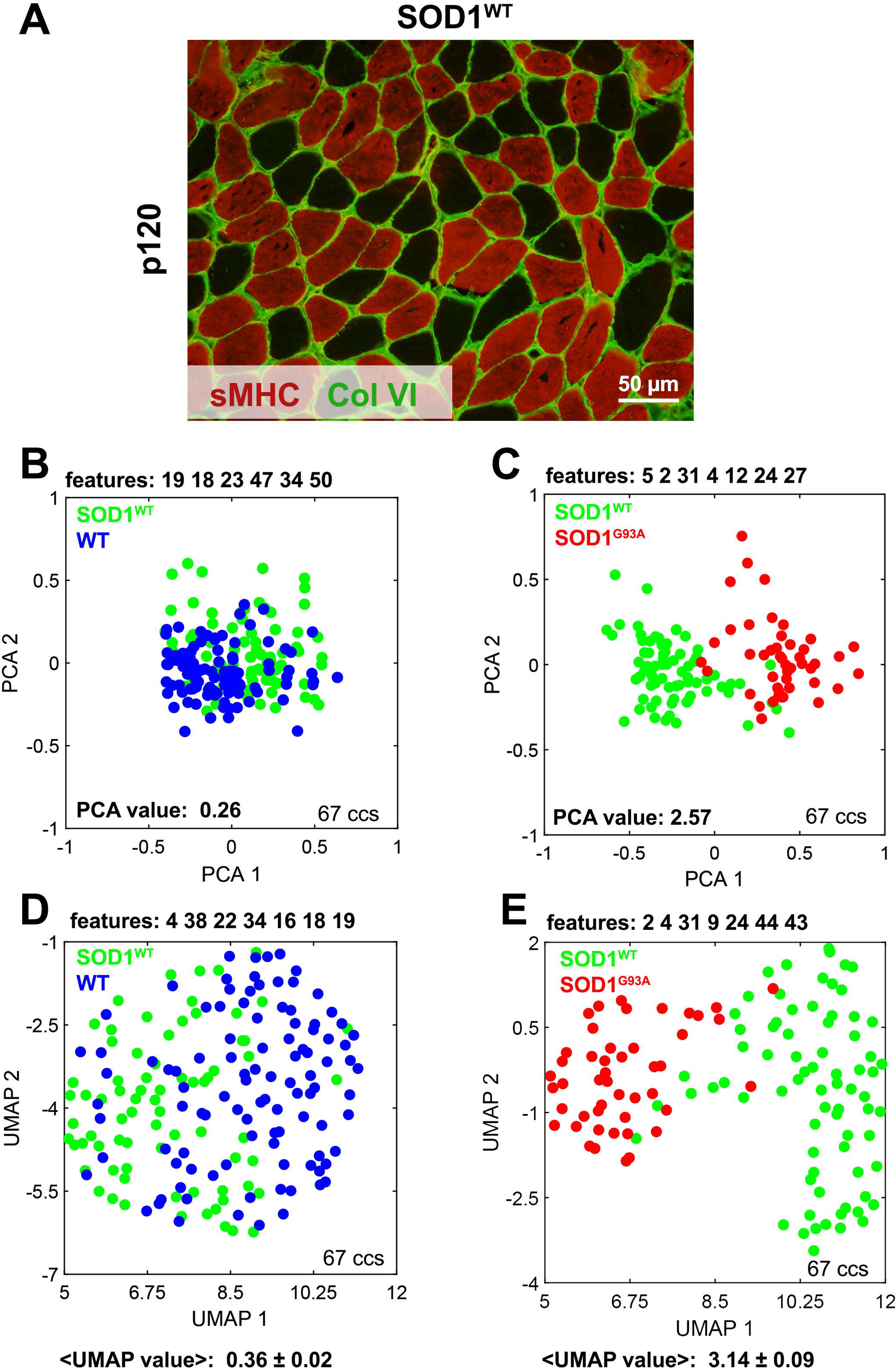
Analysis of pathological condition of SOD1^WT^ mice. Panel **A** shows an example of a p120 SOD1^WT^ soleus muscle biopsy. As in Figure 1, sMHC tags for myosin heavy chain in slow fibres, while col VI indicates the collagen VI staining. The scale bar represents 50 µm. Next, the NDICIA results comparing SOD1^WT^ vs WT (**B, D**) and SOD1^WT^ vs SOD1^G93A^ (**C, E**) at 120 days using the full set of 67 ccs are plotted. First, we handled with PCA method (**B, C**) and second with UMAP (**D, E**), Colour codes and panel organisation follow the same rules as in Figures 3 and **4**. Green dots represent SOD1^WT^ samples.

We performed several experiments to validate different conditions related to the muscle samples used in this study (**Figure S1**, **Supplementary Methods**). These computational experiments test NDICIA’s ability to obtain accurate results under unbalanced conditions, such as unequal number of images per group or sex. Additionally, we have assessed the reliability of results with small group sizes and the generality of identified features in all-vs-all comparisons for separating image subsets (**Figure S1**, **Supplementary Methods**).

### Statistics

We performed a univariate analysis to compare the values of those features extracted by NDICIA (**Table S3**). This approach allowed us to assess the individual discriminative power of each feature. We evaluated whether the feature values of these two kinds of data presented normal distribution and similar variance using the Kolmogorov-Smirnov test with the correction of Lilliefors (if n<50, we used the Shapiro-Wilk test) and two-sample F-test, respectively. If data followed a normal distribution and had similar variance, we employed a two-tailed Student’s t-test. If the data had a normal distribution but not equal variance, we employed a two-tailed Welch test to compare means from both groups. If the data did not have a normal distribution, we used the Wilcoxon test to compare medians from both groups.

### Code and data availability

The full code used in this project was developed using Matlab R2020b (Mathworks). The code is freely accessible under the GPLv3 license on https://github.com/ComplexOrganizationOfLivingMatter/MouseMuscleSOD1. Images and data necessary to reproduce this study are publicly available at https://zenodo.org/records/12795700.

## RESULTS

### Topological analysis of WT and SOD1^G93A^ mice muscles

We analysed the soleus muscle from SOD1^G93A^ and WT mice in terms of the organisation and morphology of their fibres to investigate the disease progression in this animal model for ALS. We used immunohistochemical staining in thin biopsies of these animals, enabling the identification of extracellular matrix collagen and slow muscle fibres (**Figure 1**, **Material and Methods**). Thus, we compared soleus images from SOD1^G93A^ and WT mice at different ages: postnatal day 60 (p60), 100 (p100) and 120 (p120) (**Figure 1A-F**). These subsets of images were processed using the NDICIA pipeline (**Material and Methods, Supplementary Methods).**

First, we applied the segmentation module of the NDICIA pipeline to identify the muscle fibres from the collagen network (**Figure 2A-E**, **Material and Methods**). Furthermore, fibres were classified as slow or non-slow fibres based on myosin heavy chain staining (**Figure 2B**, **2D**, **Material and Methods**). Next, we constructed a network over the segmented tissue, considering the fibres as nodes connected with their neighbouring fibres (**Figure 2F**). Second, we extracted 67 features (**Table S2**) from each image (**Supplementary Methods**). The first 14 features examined cell shape and size, while features 15-66 analysed cellular organisation and network connectivity using graph theory. The 67^th^ feature represented the proportion of slow fibres. In the final step, we applied Principal Component Analysis (PCA) combined with a feature selection algorithm to identify the most relevant combination of features for distinguishing the two groups of images (**Supplementary Methods**). The separations were scored by a PCA descriptor. A higher descriptor value indicates a greater separation between the groups [27,28] (**Supplementary Methods**). We considered that a PCA descriptor above “1” was indicative of a meaningful inter-class separation (See details in **Figure S1 and Supplementary Methods**).

To assess whether we could discriminate between WT and SOD1^G93A^ muscles and the most relevant features for these comparisons, we performed three comparisons: i) geometric features (10 ccs). ii) geometric features and slow/non-slow fibre discrimination (15 ccs). iii) using the full set of 67 features, including all geometric and network features (67 ccs). (**Material and Methods**, **Table S2**).

Using the 10 ccs subset, we first studied the latest age, p120, where the phenotypic differences between SOD1^G93A^ and WT mice were most evident. The p120 comparison revealed a strong differentiation between the two groups with a descriptor value of 2.6 (**Figure 3A, Table S3**). The features contributing most significantly to this separation included cell area variability (cc. 2); average cell convex hull (cc. 11), and standard deviation cell convex hull (cc. 12) (**Figure 3A, Table S3**). These results indicate that the mutant muscles contained cells with greater variability in area and more irregular shapes compared to WT muscles. At p100, the PCA descriptor was lower (0.74), indicating less pronounced differences between SOD1^G93A^ and WT mice (**Figure 3B, Table S3**). At this age, the clinical phenotype of the disease is present but not as severe as in p120 [15]. The selected features for this comparison included the cell minor axis (cc. 8); collagen distribution variability (cc. 14), and variability in major/minor cell axes (cc. 10) (**Figure 3B, Table S3**). Although these features indicated some differences, they were insufficient for clear separation. At p60, before ALS symptoms emerged in SOD1^G93A^ mice, the resulting PCA descriptor value (0.33) demonstrated that purely geometric features cannot differentiate the two groups at this age stage (**Figure 3C, Table S3**).

After evaluating the discriminative capability of the 10 ccs subset of features, we next evaluated the 15 ccs subset, keeping the 10 ccs and adding the slow/non-slow fibre discriminative features. In the p120 comparison, the differentiation slightly improved (from a PCA descriptor value of 2.6 to a value of 2.94, **Figure 3D, Table S3**). We captured similar features to those in the 10 ccs comparison, with the addition of the proportion of slow/non-slow fibres (cc. 67) and variability of slow fibres area (cc. 4), both of which were larger in the ALS model (**Figure 3D, Table S3**). For p100, we observed a small enhancement in the separation descriptor (from 0.74 to 0.95, **Figure 3E, Table S3**). Once again, the higher number of non-slow cells in SOD1^G93A^ compared to WT (cc. 67) and the higher variability in the non-slow fibres area were important in achieving a separation close to the threshold (separation descriptor = 1) (**Table S3**). In the p60 observation, there was an improvement in differentiation between the mouse genotypes (from PCA descriptor 0.33 to 0.88, **Figure 3F, Table S3**), although not enough for clear group separation. The inclusion of slow/non-slow fibres discrimination features, specifically the larger number of non-slow fibres (cc. 67) and the more variable area of non-slow cells (cc. 6), considerably improved the separation (**Figure 3E, Table S3**).

Finally, we applied the NDICIA protocol using the full set of 67 ccs, which included the subset of 15 ccs as well as purely graph theory features and network features that compared the geometric characteristics of the fibres neighbourhood (**Table S2**). In p120, the PCA descriptor improved further, reaching a value of 3.49 (**Figure 3G, Table S3**). Referencing the previous 15 ccs analysis (**Figure 3D, Table S3**), only two new features were added from the 67 ccs set (ccs. 24 and 27). SOD1^G93A^ showed larger variability in the neighbouring cell area (cc. 24) and a smaller average minor axis of neighbouring cells (cc. 27) compared with WT (**Figure 3G, Table S3**). For p100 mice, the PCA descriptor was found to be improved (from 0.95 to 1.12, **Figure 3H, Table S3**). Compared with the previous 15 ccs analysis (**Figure 3E, Table S3**), greater variability in neighbourhood cell area (cc. 24) and cell axes length (ccs. 26 and 30) in mutant images than in WT resulted in a clearer distinction among genotypic groups (**Figure 3H, Table S3)**. Ultimately, in the case of p60 presymptomatic mice, NDICIA was able to distinguish SOD1^G93A^ and WT images on the PCA graph using the 67 ccs set (PCA descriptor: 1.18, **Figures 1A-B** and **3I**, **Table S3**). In addition to the geometric features identified in prior analyses, the presence of some network features related to the slow fibre pattern (ccs. 20, 18, 62 and 45) played a crucial role in achieving this separation. The most significant feature in this regard was the average number of non-slow fibres that are neighbours of slow cells, with this number being smaller in SOD1^G93A^ than in WT tissues. In SOD1^G93A^, the number of neighbours of non-slow cells was more diverse than in WT (cc.18). Besides, the standard deviation of the shortest paths between non-slow cells was larger in mutants (cc. 62), suggesting that the non-slow cells were scattered and heterogeneously distributed throughout the tissue. Additionally, the average clustering coefficient of slow cells was higher in SOD1^G93A^ than in WT muscles, indicating that slow cells in mutants tended to form larger clusters (cc. 45).

Importantly, we compared the values of the selected features from the WT and SOD1^G93A^ samples (**Figure 3G-I**) using univariate statistical analysis (**Table S3**, **Material and Methods**). Despite some individual features showing significant differences in their values, none were shared across the p60, p100 and p120 comparisons (**Table S3**). This further emphasises the importance of NDICIA analysis, which integrates contributions from multiple features to achieve accurate differentiation between groups, enabling a comprehensive and objective analysis of disease progression in ALS.

### Uniform Manifold Approximation and Projection (UMAP) method to upgrade the dimensionality reduction module

PCA is a well-established method for dimensionality reduction through linear transformation, typically projecting data into a two-dimensional plane. To incorporate non-linear transformation, we upgraded NDICIA by integrating the Uniform Manifold Approximation and Projection (UMAP) algorithm [26] (**Material and Methods**). This integration allowed us to explore novel factors contributing to the separation of the samples. The UMAP algorithm was run 100 times per comparison to mitigate stochasticity effects, capturing the most frequently repeated features and computing the average UMAP descriptor (**Material and Methods**). The NDICIA workflow remained unchanged, except for replacing PCA with UMAP. Therefore, we tested the same groups of characteristics: i) 10 ccs, ii) 15 ccs and iii) the complete set of features, 67 ccs:

In the geometric evaluation (10 ccs), UMAP yielded descriptors very similar to the PCA method (**Table S3**). In p120, a high average UMAP descriptor (average UMAP descriptor: 2.44±0.87) differentiated SOD1^G93A^ from WT images (**Figure 4A, Table S3**). The more irregular and variable cell shape (ccs. 2, 8, 10, 11 and 12) and the larger heterogeneity of collagen distribution (cc. 14) in mutants than in WT were the features driving this separation. However, in p100 (average UMAP descriptor: 0.63±0.03) and p60 (average UMAP descriptor: 0.37±0.04), clear separation was not achieved (**Figure 4B-C, Table S3**). The inclusion of slow and non-slow fibre discrimination to geometric features (15 ccs) improved separation across all age-stages separations. At p120, a larger standard deviation of slow and non-slow fibres area (ccs. 4 and 6) in SOD1^G93A^ than in healthy mice enhanced the dissociation (average UMAP descriptor: 3.25±0.15) (**Figure 4D, Table S3**). In p100, the greater variance in non-slow fibre area (cc. 6) and the higher proportion of slow/non-slow fibres (cc. 67) in mutants than in WT, as in PCA, raised the UMAP value close to 1 (average UMAP descriptor: 0.94±0.04) (**Figure 4E, Table S3**). Similarly, at p60, the larger proportion of slow fibres (cc. 67) and variability in non-slow fibre areas (cc. 6) improved separation close to 1 (average UMAP descriptor: 0.95±0.08) (**Figure 4F, Table S3**). Finally, using the whole set of 67 features, the separation of all age groups was slightly enhanced using UMAP compared to the PCA method (**Figure 4G-I**, **Table S3**). p120 comparison (average UMAP descriptor: 3.95±0.21) highlighted features related to heterogeneous fibre shape and size, such as standard deviation in cell area (ccs. 2, 4), convex hull (ccs. 11, 12), and neighbour relationships in fibre geometry (ccs. 23, 27, 28) (**Figure 4G**, **Table S3**). All these features pointed to more heterogeneous fibre shape and size in the mutant than in WT mice, which is a similar result to that observed in PCA (**Table S3**). In p100 (average UMAP descriptor: 1.26±0.03, **Figure 4H, Table S3**), three of the selected features (ccs. 14, 6, 30) were also seen in the PCA analysis. In addition, UMAP revealed greater variation in fibre shape (cc. 31) and highlighted network features related to fibre type (ccs. 20, 58, 45), which suggests that slow cells in SOD1^G93A^ are more clustered than in WT. In the p60 comparison (average UMAP descriptor: 1.30 ± 0.10, **Figure 4I, Table S3**), three of the selected features (ccs. 20, 18, 62) matched those from the PCA analysis. As seen in the p100 analysis, ccs. 6 and 31 emerged due to the greater variability in non-slow cell size (cc. 6) and the more diverse convex shapes of neighbouring cells (cc. 31) in the ALS model compared to WT. Also, cc. 8 was selected because mutant fibres had a larger minor axis. Additionally, cc. 22 was selected probably because non-slow fibre arrangement is more scattered in SOD1^G93A^ tissue (less clustering of non-slow cells) than in WT, which are more grouped.

Importantly, the univariate statistical analysis of the features appearing to differentiate mutant and WT at different age stages showed mostly significant differences when comparing SOD1^G93A^ and WT features values (*p*-value<0.05 *, **Table S3**, **Material and Methods**). Moreover, as in PCA, none of them was both selected and significant in the three age comparisons (**Table S3**). However, in general terms, the number of individual significant differences after using UMAP was equal to or better than using PCA.

### NDICIA to study the role of SOD1^WT^ mice as an ALS model

The previous results have demonstrated that NDICIA can find differences between samples from WT and affected mice at all age stages. To confirm the biological relevance of the differences found while rejecting an overfitting scenario, we introduced a third animal model: The SOD1 wild-type (SOD1^WT^) genotype, which overexpresses the non-mutant human SOD1 protein.

Some studies propose that SOD1^WT^ protein may act as a causal factor in cellular senescence [29] or share pathological pathways with mutant SOD1 [30–34]. However, the role of SOD1^WT^ models in ALS remains a topic of longstanding scientific debate [32,35], and its significance has yet to be fully established. We focused on the p120 stage, where the separation between WT and SOD1^G93A^ samples was clearest, and compared both with SOD1^WT^ images (**Figures 1E-F** and **5A**). First, using the PCA strategy, in the case of the comparison between WT and SOD1^WT^ samples, the PCA value was very small indicating that both sets of images were similar (PCA descriptor: 0.26, **Figure 5B, Table S3**). However, when we compared SOD1^WT^ and SOD1^G93A^, we observed a clear separation between the samples displaying a high PCA descriptor (PCA descriptor: 2.57, **Figure 5C, Table S3**). Interestingly, 5 out of the 7 features (ccs. 2, 4, 12, 24 and 27) were also present in the PCA comparison between p120 SOD1^G93A^ and WT samples (**Figure 3A, Table S3**). The results were consistent when we applied the UMAP approach. In the case of WT vs SOD1^WT^, it was not possible to find a separation between the groups (average UMAP descriptor: 0.36±0.02, **Figure 5D, Table S3**), whereas SOD1^G93A^ and SOD1^WT^ were differentiated (average UMAP descriptor: 3.14±0.09, **Figure 5E, Table S3**). Both sets of results suggested that SOD1^WT^ samples are very similar to WT in terms of muscle fibre shape, size and organisation, and drastically different from SOD1^G93A^.

## DISCUSSION

As of now, a conclusive ALS diagnostic test or biomarker remains elusive. Currently, ALS diagnosis relies mainly on physician observation of patient signs, along with electromyography [36] and a battery of tests to rule out other diseases [37,38]. These clinical data allow physicians to determine different evidence levels for suffering from ALS (suspected, possible, probable and definitive diagnosis). The importance of identifying biomarkers to validate new treatments is evident. In this work, we study the SOD1^G93A^ mouse, a well-known murine model in ALS preclinical research [39]. By using NDICIA, we quantify and evaluate differences between mutant and healthy mouse muscles, with the ultimate goal of identifying potential ALS image biomarkers. Here, we show how our computational approach unveils several neurogenic indicators in mouse presymptomatic stages.

Our robust multivariate analysis using the NDICIA software protocol [24] has been successful in reporting differences in muscle organisation in human samples from different muscle groups [23], and separating healthy, neurogenic disorder and muscle dystrophy samples [22]. Applying NDICIA to our p100 and p120 muscle samples, after ALS symptoms presumably emerged, we found clear separations of WT and mutant phenotypes in terms of PCA value, with the separation becoming more pronounced as the illness progresses. We found that the main properties driving the separation between both groups were related to the fibres’ geometry, which was more homogeneous in size and shape in WT than in SOD1^G93A^, revealing the pattern of myofibre neurogenic atrophy in affected tissue. Additionally, the higher quantity and heterogeneous distribution of collagen along the SOD1^G93A^ tissue were also important in the analysis. These results are evident simply by comparing WT and SOD1^G93A^ biopsies through visual inspection. The graph theory features associated with the type of fibre (slow/non-slow) did not play an important role in the p100 and p120 separations. On the other hand, the separation between WT and SOD1^G93A^ at the presymptomatic stage (p60) was primarily based on network properties related to fibre-type discrimination. The results suggest the presence of larger clusters of slow fibres in SOD1^G93A^ muscle tissues than in WT, where non-slow fibres are scattered throughout the tissue. Although the proportion of slow/non-slow fibres is always larger in mutants at any stage (p60, p100 or p120), and this could bias the network results, it is only at p60 that these kinds of properties enhance the separation. These findings could be associated with the early changes in neuromuscular transmission detected in [40], long before the onset of motor symptoms. Additionally, the higher number of slow fibres in SOD1^G93A^ could be related to the well-known preferential denervation of fast fibres that, in a second step, is partially compensated by fast-to-slow fibre type transitions [41].

The ability of NDICIA to detect differences in samples that appear similar upon visual inspection might suggest that it forces the separation of any two groups of images. However, our results with the SOD1^WT^ mice reject this hypothesis. We were unable to find differences between WT and SOD1^WT^ samples at p120 with the whole set of 67 features. Prior studies suggest misfolded SOD1 aggregates or shared pathways with mutant SOD1 may link SOD1^WT^ to ALS [29–31,42,43], challenging its use as a control. Our results indicate that SOD1^WT^ is similar to WT in terms of muscle fibre characteristics, contradicting the idea that SOD1^WT^ overexpression plays a pathogenic role in ALS and reinforcing its characterisation as a healthy model.

We acknowledge that our approach, in general, and our graph theory analysis, in particular, has certain limitations that may raise concerns about the accuracy of the results. These challenges include the presence of cryoartefacts commonly found in muscle biopsy preparations and the small or unbalanced sample sizes. Regarding the relevance of the graph theory features, while constrained by the low number of fibres contained within an image, they are sufficient to capture organisational patterns at p60 (i.e. finding a specific and presymptomatic progressive pattern of fast-to-slow fibre-type transitions, resulting in grouping slow cells in a restricted way). It is possible that additional markers, such as fast-myosin, could provide further features for graph theory analysis, enhancing the differentiation between mutant and control samples. However, we consider one of NDICIA’s strengths is its ability to work using only two primary antibodies. The use of images obtained by basic and routine protocols makes it potentially applicable to clinical labs. Regarding the number of images, our validation experiments and previous studies demonstrate that small or unbalanced sample sizes do not compromise the reliability of NDICIA’s outcomes (**Figure S1, Supplementary Methods**) [22,23]. Furthermore, NDICIA’s robustness allows it to handle heterogeneous image classes while consistently delivering reliable and generalisable results. We conclude that, while limitations exist, they are effectively mitigated by NDICIA’s strengths, making it a trustworthy tool for muscle analysis. Related to this aspect, we think that the ability of NDICIA to process images that are easy to obtain and that may contain little artefacts enhances its potential for identifying ALS image biomarkers, emphasising the significance of network features in early diagnosis.

We have been actively working to enhance NDICIA’s performance. Here, we present an upgrade of NDICIA achieved through the integration of UMAP, a nonlinear dimensionality reduction method, into the pipeline. UMAP is a powerful tool for visualising complex data, offering greater sensitivity than previous methods [44–46]. Thus, to get possible better separation and different combinations of selected features, taking advantage of the non-linear relationships considered by UMAP, we applied it to the same WT-SOD1^G93A^ comparisons previously analysed with PCA. A limitation of UMAP is its stochastic nature; therefore, we performed 100 repeated simulations per experiment to ensure consistent average results. When analyses were carried out after considering the whole set of features (67 ccs), all the NDICIA results slightly improved. Consequently, UMAP supports the main conclusions derived from PCA analyses. In addition, UMAP’s performance has been validated to be used within NDICIA for application to similar studies. Building on the current capabilities of NDICIA, our plans include implementing AI-based algorithms for image segmentation and classification, leveraging the data generated throughout this work as a training dataset.

In summary, our study highlights the utility of NDICIA to preclinical studies, identifying neurogenic signatures in ALS-like muscle as a potential biomarker, even when the disease is phenotypically imperceptible. This early pathological detection could drive the specific preclinical or even clinical trials, bringing forward the onset of a potential treatment before muscle degeneration becomes consequential. Given these findings, together with the previous application of NDICIA to patient samples [22,23], our tool holds promise for future clinical application. However, further investigation and validation using patient images will be essential to fully realise its potential.

## AUTHOR CONTRIBUTIONS

A.M.C. worked in the sample preparation and imaging. V.N. supervised the sample preparation and contributed to the design of the figures. P.G.-G. carried out the computational experiments and designed the figures and tables. L.M.E. and C.P. conceived the study and supervised the experiments. P. G.-G., L.M.E. and C.P. wrote the paper with input from all authors. All authors participated in the interpretation of the results, discussions, and development of the project.

## ETHICAL APPROVAL

The authors declare that ethical approval for this project was granted by the local research ethics committee.

## ABBREVIATIONS

ALS: Amyotrophic Lateral Sclerosis
ccs: Characteristics
NDICIA: Neuromuscular DIseases Computerized Image Analysis
SOD1: Superoxide dismutase 1
WT: Wild-type
p60, p100 and p120: Postnatal day 60, 100 and 120 subsequently
PCA: Principal Components Analysis
UMAP: Uniform Manifold Approximation and Projection

