## Supplementary Methods for "Computational analysis of SOD1-G93A mouse muscle biomarkers for comprehensive assessment of ALS progression"

**LEGENDS OF SUPPLEMENTARY FILES**

**Figure S1. Validation experiments.** A) Distribution of PCA descriptor values after 1000 NDICIA PCA realizations comparing randomly selected 25-vs-25 images subsets from p120 WT and SOD1^G93A^ (green bars). The distribution of PCA descriptor values after random class shuffling of these comparisons (grey bars) is also shown. Class A consists of 12 WT images and 13 SOD1^G93A^ images while class B contains 13 WT and 12 SOD1^G93A^ images. The red dashed line indicates inter-class separation threshold, and the black arrow marks the PCA separation descriptor from the p120 all-vs-all comparison (**Figure 3G**). B) Most frequently identified features across 1000 realizations of “25 WT vs. 25 SOD1^G93A^” comparisons. The features highlighted in italic bold green correspond to those also selected in **Figure 3G**. Green bars represent the top 10 selected features, while grey bars indicate features ranked 11^th^ to 20^th^. C) PCA descriptor values after running NDICIA PCA using only the seven features preselected in **Figure 3G**. D-L) NDICIA PCA, sex analysis. D) p60, only females. E) p100, only females. F) p120, only females. G) p100, only males. H) p120, only males. I) p100, WT males vs. SOD1^G93A^ females. J) p100, WT females vs. SOD1^G93A^ males. K) p120, WT males vs. SOD1^G93A^ females. L) p120, WT females vs. SOD1^G93A^ males. Note that most p60 samples (45 in 46) come from female mice; therefore, comparisons with males are not possible. All NDICIA PCA experiments were performed using 67 ccs.

**Table S1. Image dataset.** Our dataset comprises 386 images collected from 61 mice. In the p60 group, we collected 45 images from 7 female mice and 1 image from 1 male; 27 images from 5 WT mice and 19 images from 3 SOD1^G93A^ mice. For the p100 group, 67 images were acquired from 8 males and 48 images from 10 females; 78 images from 13 WT mice and 37 images from 5 SOD1^G93A^ mice. In the p120 group, 111 images were obtained from 18 male mice and 114 from 17 females; 103 images from 11 WT mice, 47 images from 13 SOD1^G93A^ mice, and 75 images from 11 SOD1^WT^ mice.

Logistical constraints, including funding limitations and reduced technical staffing, impacted on our ability to sustain the mouse colony and collect large datasets for every group. These challenges resulted in differences in the number of images across age-group comparisons. Validation experiments demonstrated that a relatively small number of images (~20-25 per group, similar to p60 comparison) is sufficient to extract consistent image biomarkers (**Figure S1**). These validation experiments yielded reliable conclusions comparable to those obtained from larger datasets (**Figures 3** and **S1**). Additionally, group differences were confirmed to be independent of the mouse sex (**Figure S1**).

**Table S2. Set of features analysed using NDICIA.** The table shows the names and descriptions of 67 characteristics analysed in this article. The characteristics can be classified as geometric (1-14), network (15-66) and the proportion of slow cells (67). The bold names mark the properties related to slow or non-slow cell type discrimination.

**Table S3. Quantitative and statistical results after NDICIA application.** This table is organized in three sheets, 67 ccs, 15 ccs and 10 ccs. Each one collects all NDICIA results for each comparison developed along the manuscripts using as source the subset of 67, 15 or 10 ccs: PCA descriptor; selected features; absolute weights of these features in the PCA graph discrimination; UMAP descriptor; average and standard deviation of the features values in each compared group, and an individualized statistical analysis to explore if the selected features present significant difference when comparing the different groups of images. The statistical tests were selected using a protocol depending on the data characteristics (**Material and Methods**), and the goodness of the statistical difference was evaluated by three levels of *p*-value: <0.05*, <0.01**, 0.005***.

**SUPPLEMENTARY METHODS**

**Validation experiments**

To validate the diverse sample sizes per group and assess the reliability of our results (**Figure 3**), we conducted additional experiments supporting our approach and conclusions. These experiments also confirmed PCA descriptor > 1 as a robust threshold for distinguishing image groups. All validation experiments were performed using the NDICIA PCA pipeline, as it ensures deterministic results. The use of just this methodology for the validation experiments is supported by the results obtained with NDICIA UMAP as they were consistent with those from NDICIA PCA (**Figures 3** and **4**).

We first evaluated whether our method reliably compared small datasets and produced results comparable to those from larger datasets using the full set of 67 features. For p120, where group differences were most pronounced (**Figure 3H**), we designed an unbiased approach to probe the NDICIA pipeline. This approach involved 1000 NDICIA PCA realizations, each randomly selecting 25 WT and 25 SOD1^G93A^ images. The PCA descriptor was consistently greater than 1 (**Figure S1A**). Conversely, when shuffling the classes to create groups with approximately equal proportions of WT and SOD1^G93A^ images (e.g., 12 and 13 per condition), the descriptor was always below 1 (**Figure S1A**). These findings indicate that NDICIA does not separate groups lacking genotypic differences.

Additionally, the most frequently identified features across the 1000 iterations (**Figure S1B**) matched most of those identified in the overall WT vs. SOD1^G93A^ p120 comparison (**Figure 3G**). 6 out of 7 features from **Figure 3G** were found in the top 10 of selected features across the 1000 “25 WT vs. 25 SOD1^G93A^” realizations (**Figure S1B**). To further validate the generalisability of these features, we performed 1000 NDICIA PCA comparisons using only the seven features identified in **Figure 3G**. These features consistently separated the 25-vs-25 images groups across all realizations (**Figure S1C**).

Finally, we tested NDICIA exclusively on female images (**Figure S1D-F**), mirroring the p60 analysis where most images were from female mice (**Table S1**, **Figure 3I**). Further comparisons at p100 and p120 groups, including [male vs male], [female vs female] and [male vs female] analyses, confirmed sex independence of our results (**Figure S1G-L**).

Together with previous random comparisons accounting for sex and tissue region variability, these results demonstrate that NDICIA provides reliable outcomes regardless of tissue sampling region, sex, or sample size per class.

**NDICIA modules: Preprocessing and segmentation**

Segmentation is the first step in the image processing protocol. At this stage, the collagen network and muscle fibres must be clearly differentiated (**Figure 2E**). This process is semi-automatic and relies on morphological operations implemented in MATLAB R2020b (MathWorks), following the methodology described by [1]. The procedure begins with the separation of colour channels in RGB images obtained through microscopy. The green channel, representing the extracellular matrix primarily composed of collagen fibres (**Figure 2C**) is pivotal for segmenting the muscle fibres by delineating their contours, while the red channel highlights slow-twitch fibre staining.

After selecting the green channel, the intensity map is auto-adjusted to enhance contrast, enabling better differentiation between muscle fibres and the extracellular matrix. A segmentation method based on h-minima and h-maxima transformations is then applied, as outlined by [1]. This type of segmentation has been successfully applied in several medical enforcements [1–3]. Through this approach we transform the green channel of the image using different h-values. For each h-value, we obtain a range of intensities where pixels representing the extracellular matrix are connected. Overlapping the pixels from individual transformations creates a complete connective tissue mesh, where the “holes” correspond to muscle fibres.

Subsequently, noise and artefacts are removed by comparing the areas of individual cells. Morphological operations, including opening, closing, hole filling, and watershed transformation, are performed to fill small gaps, compact structures, and separate joined fibres. Finally, manual curation is carried out by an expert to refine the segmentation. The software used was Adobe Photoshop CS6.

**NDICIA modules: Slow fibre detection**

The detection and classification of muscle fibres into slow and non-slow are conducted after segmentation. Since the red channel (R) specifically marks slow fibres, a muscle fibre is classified as slow if it meets at least one of the following criteria: 1) The average intensity value within the fibre is higher than a quarter of the maximum intensity value in the red channel image. 2) At least 15% of all intensity values within the fibre area exceed half of the maximum intensity value of the red channel image. These criteria prevent the misclassification of non-slow fibres due to isolated intensity peaks caused by image artefacts.

Muscle fibres that do not meet these criteria are classified as non-slow fibres. After this automatic process we ensure the classification with a supervised review of the whole set of images by using a custom Matlab script.

**NDICIA modules: Features extraction**

Once muscle fibre contours are accurately segmented and fibre type is classified, various tissue properties were extracted and quantified (**Figure 2F**, **Table S2**):

Geometric features. These features include the area, the minor and major axis lengths, and the geometric shape of muscle fibres (ccs. 1-12), along with the amount of collagen surrounding them (ccs. 13-14). The collagen region associated with a specific fibre is calculated using the Voronoi algorithm [4], with the segmented fibres serving as Voronoi seeds. This approach defines the equidistant boundaries between muscle fibres and assigns the collagen enclosed within these boundaries to the corresponding fibre.

Network features. These features are extracted based on the connectivity between muscle fibres. Connections between fibres are defined based on the shared boundaries identified using the Voronoi algorithm. Two fibres sharing a boundary are considered neighbours. From each neighbourhood, relational features are computed, including the number of neighbouring fibres for each fibre and the ratios of a fibre’s geometric properties (e.g., area, major axis, minor axis, convexity, or surrounding collagen) compared to those of its neighbours (ccs. 15-34, **Table S2**). Additionally, by modelling the tissue as a network, graph theory metrics such as eccentricity, clustering coefficient, strength, betweenness centrality, and shortest path length are computed (ccs. 35-66, **Table S2**). Lastly, the proportion of slow and non-slow fibres in the muscle sample is calculated (cc. 67, **Table S2**).

**NDICIA modules: Feature selection**

The final step of NDICIA involves a feature selection module, which follows a sequential process: i) Execute the dimensionality reduction method (see “NDICIA modules: Reduction of dimensionality”) for every pair of features, retaining the 10 pairs with the highest separation descriptor (see “NDICIA modules: Descriptor of separation”). ii) Expand each of the top 10 feature pairs into trios by testing all possible combinations, retaining the 5 trios with the highest separation descriptor. iii) For the top 50 trios, identify and retain the two quartets with the highest separation descriptor from each trio, resulting in up to 100 quartets. iv) Incrementally add one feature at a time to the quartets, selecting the feature that most improves the separation descriptor. v) Repeat the incremental feature addition until the combinations include seven features or the separation descriptor ceases to improve.

Feature importance (or weight) is assessed based on the absolute values of their eigenvectors leading to transformation into 2D projections. In NDICIA PCA plots (**Figures 3** and **5B-C**), features are ranked above the graphs in descending order of weight. In contrast, for NDICIA UMAP plots (**Figures 4** and **5D-E**), features are ranked based on their frequency of appearance across 100 realizations, displayed from left to right.

**NDICIA modules: Reduction of dimensionality**

In NDICIA, dimensionality reduction was initially performed using the PCA algorithm, a classical method that assumes data can be expressed as a linear combination. PCA transforms the data into two principal components, typically represented as axes capturing the largest variance within the dataset.

Aiming to upgrade the dimensionality reduction capabilities of NDICIA, PCA was replaced with UMAP, a novel manifold learning technique that effectively handles non-linear relationships [5]. UMAP is a stochastic method that requires the tuning of several hyperparameters, including:

n_neighbours: Determines the local neighbourhood size used for manifold approximation. Larger values provide a more global view of the manifold, while smaller values preserve local structures. Range: 0 – 100,

min_dist: Sets the minimum effective distance between embedded points. Smaller values result in more clustered embeddings, while larger values create a more even distribution. Range: 0 - 0.99.

n_components: Specifies the target dimensionality of the embedding space.

metric: Defines the distance metric for high-dimensional space.

For this study, the UMAP MATLAB implementation (v1.3.4) [6] was employed using the default hyperparameter settings: ‘n_neighbours’ = 30, ‘min_dist’ = 0.3, ‘n_components’ = 2 and ‘metric’ = euclidean. These values were selected to achieve a balance between capturing local and global variations in the embedding.

To integrate UMAP with the feature selection method (see “NDICIA modules: Feature selection”) and ensure reliability despite inherent stochasticity, the feature selection pipeline was repeated 100 times. Each realization produced a combination of features optimally separating two image groups. The most frequently occurring features across the 100 realizations were selected. If the typical feature combination length was six, only the six most frequently repeated features were retained.

Additionally, the mean and standard deviation of the UMAP descriptor (see “NDICIA modules: Descriptor of separation”) were calculated by averaging all UMAP descriptors from each realization. A representative visualization was selected based on the realization with the UMAP descriptor closest to the average UMAP value (**Figures 4** and **5D-E**).

**NDICIA modules: Descriptor of separation**

After reducing dimensionality using PCA or UMAP algorithms (see in “NDICIA modules: Reduction of dimensionality”), the transformed data allow the spatial distribution of the “points” (images) to be analysed in a two-dimensional graph. The degree of separation between the two image groups is then evaluated using a variant of the Calinski-Harabasz descriptor [7], as detailed in [8]. This measure is referred to throughout the manuscript as PCA or UMAP descriptor, depending on the algorithm used. This descriptor is mathematically described as the measurement of the scatter between class means ($B$, how far apart the class centers are) in relation with the scatter within class ($W$, how spread out the points are around their respective class means) defined by $descriptor=trace\left( \frac{B}{W} \right).$

$$W=\sum_{i=1}^{k} \sum_{l=1}^{N_{i}} \left( x_{i}\left( l \right)-\bar{x_{i}} \right)\left( x_{i}\left( l \right)-\bar{x_{i}} \right)^{T}$$

$$B=\sum_{i=1}^{k} N_{i}\left( \bar{x_{l}}-\bar{x} \right)\left( \bar{x_{l}}-\bar{x} \right)^{T}$$

Where $X=\{x(1), \ldots, x(N)\}$ is a set of $N$ data objects and a partition of these data into $k$ mutually disjoint clusters, $N_{i}$ is the number of objects assigned to a $i_{th}$ cluster, $x_{i}(l)$ is the $l_{th}$ object assigned to that cluster, $\bar{x_{l}}$ is the n-dimensional vector of sample means within that cluster (cluster centroid) and $\bar{x}$ is the n-dimensional vector of overall sample means (data centroid). As such, the within-group and between-group matrices sum up to the scatter matrix of the data set. Consequently, compact and separated clusters are expected to have small values of $W$ and large values of $B$. Hence, the better the data partition the greater the value of the ratio between $B$ and $W$ [7].

When $descriptor<1$, intra-class variance leads over inter-class variance, and results into overlapping clusters, while when $descriptor>1$, inter-class variance dominates resulting in distinguishable and separate clusters. To challenge this assumption, we run some validation experiments that evidence $PCA_{descriptor}=1$ is a good threshold for class separation (**Figure S1**).
