## Supplementary figures and images for "Computational analysis of SOD1-G93A mouse muscle biomarkers for comprehensive assessment of ALS progression"

### Figure S1

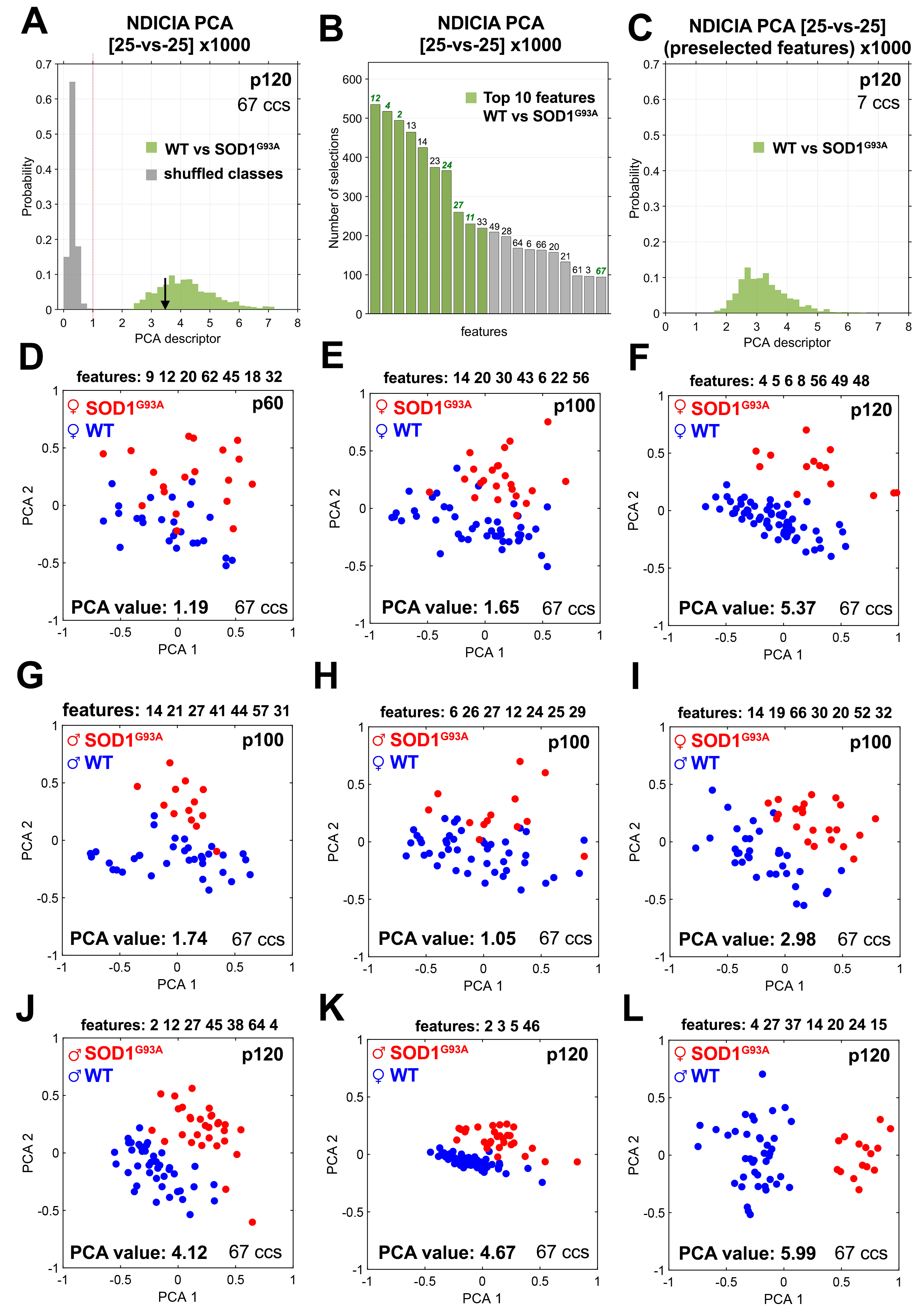
